# Physiological responses to drought stress and recovery reflect differences in leaf function and anatomy among grass lineages

**DOI:** 10.1101/2022.07.30.502130

**Authors:** Seton Bachle, Marissa Zaricor, Daniel Griffith, Fan Qui, Christopher J Still, Mark C Ungerer, Jesse B Nippert

**Affiliations:** Division of Biology, Kansas State University, Manhattan, Kansas, United States of America; Department of Plant Biology, Michigan State University, Lansing, Michigan, United States of America; Department of Forest Ecosystem and Society, Oregon State University, Corvallis, Oregon, United States of America

**Keywords:** Poaceae, Drought Response, Plant Functional Traits, Anatomy, Ecophysiology, Leaf Economic Spectrum

## Abstract

Grasses are cosmopolitan, existing in many biome and climate types from xeric to tropical. Traits that control physiological responses to drought vary strongly among grass lineages, suggesting that tolerance strategies may differ with evolutionary history. Here, we withheld water from 12 species representing six tribes of grasses to compare how species respond to drought in different grass lineages. We measured physiological, morphological, and anatomical traits. Dominant lineages from tropical savannas, like Andropogoneae, tolerated drought due to above and belowground morphological traits (specific leaf area and root length, *SLA* and *SRL*), while temperate grasses in this study utilized conservative leaf physiology (gas exchange) and anatomy traits. Increased intrinsic water-use efficiency coincided with a larger number of stomata, resulting in greater water loss (with inherently greater carbon gain) and increased drought sensitivity. Inherent leaf and root economic strategies impacting drought response were observed in all species, resulting in either high *SLA* or *SRL*, but not both. Our results indicate that grasses subjected to severe drought were influenced by anatomical traits (e.g., number of stomata and xylem area) and similar within lineages. In addition, grasses recovered at least 50% of physiological functioning across all lineages and 92% within Andropogoneae species, illustrating how drought can influence functional responses across diverse grass lineages.

## INTRODUCTION

Grasslands play a major role in regional carbon sequestration and water cycling because grasses invest in extensive rooting systems and storage organs [1,2]. Carbon dynamics are highly influenced by water availability in grassland systems, evident in drought years that result in decreased productivity [3–5]. Grasslands experiencing extreme droughts can have reduced physiological functioning [6,7], increased invasibility from non-native species [8], disruption of fire intervals [9], and loss of ecosystem functioning (i.e., productivity & species composition) [10–12]. While many grass species in grassland ecosystems have evolved in the context of an inherently variable climate, future climate projections emphasize large shifts in water availability, resulting in extreme drought and deluge events within the coming century [7,12–15]. While it is widely accepted that grasslands will vary in drought response (ability to withstand shifts from equilibrium) and drought recovery (ability to regain equilibrium), modifications in precipitation seasonality and amount will have sizable and diverse impacts on ecosystem function [16–18].

Biophysical factors determining drought sensitivity in individual plant species include precipitation and temperature variability [19], while biotic factors such as plant productivity, species richness [20], and potentially dominant species with associated functional traits, also play an important role in observable drought responses [21]. Furthermore, the history of different drought exposure in plant lineages is likely to frame future drought responses within those lineages. For example, lineages of plant species from arid and semi-arid regions have functional traits (narrow leaves, strict stomatal regulation, absorptive rooting systems) that allow them to acquire and conserve water [22], whereas lineages from wetter and warmer regions may have wider leaves and altered stomatal traits that result in distinct water-use strategies [23,24]. These evolutionary tradeoffs have shaped functional differences across lineages and directly impact ecological dynamics [25]. However, the extent of such evolutionary tradeoffs has not been utilized to identify lineage-specific trait responses to extreme drought conditions. Even more uncommon are investigations that combine physiology, anatomy, morphology, and structural data from grass species spanning several Poaceae tribes.

Large interannual variation in precipitation is a feature of many grassland ecosystems and, in combination with CO_2_ and temperature, has played a major role in the evolution and biogeographic history of major grass lineages [26–28]. Importantly, the varying evolutionary histories of grasslands have driven the evolution of different functional traits across the Poaceae phylogeny, likely accounting for differences in drought responses [12,22]. For example, leaf-level anatomical trait variation and convergent evolution has resulted in spatially separated photosynthetic tissues allowing for C_4_ photosynthesis, which is heavily expressed in Poaceae, and provides a physiological advantage that increases carbon assimilation while reducing water loss via stomatal regulation [29,30]. While it is recognized that C_4_ species are not inherently more drought tolerant than C_3_ species [12,31,32], there is evidence that increased *WUE* (water-use efficiency), inherent to C_4_ species, can be advantageous when water is limiting [33–35]. For example, native species in the arid American southwest, have the ability to initially tolerate the negative consequences of drought by maintaining physiological functioning for prolonged periods of time [36,37]. The ability of some species to maintain physiological functioning despite drying soils may be due to increased cuticle thickness, decreased stomatal size and densities, less negative turgor loss point, and more conservative growth strategies (specific leaf area, *SLA*; specific root length, *SRL*) [22,38–40]. Alternatively, the production of less metabolically costly leaves and roots (higher carbon to nitrogen ratio) and tight stomatal regulation is associated with the ability to avoid desiccation and quickly recover once drought breaks [41–43]. The ability to quickly resume pre-drought physiological function via rapid recovery may or may not be associated with the ability to tolerate drought in the first place [16,17]. Thus, drought responses are determined via morphological, physiological, and anatomical traits; all of which have been shaped from evolutionary histories.

During the evolutionary development of Poaceae, separate lineages have evolved different suites of traits, including fairly different water use strategies [26,44]. For instance, the two most abundant monophyletic groups of C_4_ grasses - Andropogoneae (water spenders) and Chloridoideae (water savers) - vary in water-use strategies because of distinct biogeographic histories [25,45,46]. Species in these lineages occupy warmer climates but vary in global distribution as a function of precipitation availability: high in Andropogoneae and low in Cynodonteae [47–49]. There are many characteristics impacting water-use and drought response associated with this ecological sorting, and they include morphological, physiological, and anatomical traits. Morphological strategies and traits associated with water relations include the production of fine roots to increase water absorption [50–52], leaf rolling to decrease irradiance [53], and variations in growth form (caespitose and rhizomatous) [54,55]. These traits are often related in terms of economics, reflecting plant investment of carbon and nitrogen in both leaf and root structures [56]. More specifically, these morphological traits are framed by underlying structures at the anatomical level in leaf and root tissues [57]. Anatomical leaf traits within and across families in Poaceae also have been observed to influence physiological responses most often associated with hydraulics (xylem lumen area/diameter; resistance to cavitation) [58–61]. However, the aforementioned physiological, morphological, and anatomical traits may not convey equal benefits in drought response or recovery across and within Poaceae lineages. For these reasons, it is increasingly important to understand how diverse lineages of grass species that vary in climate niches and evolutionary histories will respond to extreme drought conditions

Here, we conducted a robust assessment of physiological and anatomical traits from multiple grass lineages in response to and following recovery from soil drying. The species under investigation were selected based on divergent drought responses within lineages. We performed a dry-down experiment to impose severe drought on 12 species of grasses across six tribes within the Poaceae lineage. We withheld water in order to assess various physiological, morphological, and anatomical trait responses to drought, as well as (above and belowground) productivity data, to capture both drought response and recovery. We hypothesized that: (1) species within tribes will exhibit a similar response to drought sensitivity (duration in drought), based on similar evolutionary histories and drought traits specific to withstanding long periods of low water availability; (2) species within tribes will also exhibit similar responses in drought recovery, based on shared evolutionary histories and functional traits that serve to quickly utilize resources when available; and (3) leaf-level anatomical traits would best describe the species (within and across tribes) response to, and recovery from drought due to the constraints of structures that influence water transport and availability.

## MATERIALS AND METHODS

Twelve grass species from six tribes were grown from seeds obtained from the USDA Germplasm Resources Information Network or locally sourced from the Konza Prairie Biological Station. Species include: *Paspalum juergensii, Paspalum notatum, Festuca ovina, Panicum virgatum, Setaria viridis, Urochloa ruziziensis, Andropogon gerardii, Sorghastrum nutans, Danthonia spicata, Rytidosperma semiannulare*, *Bouteloua dactyloides*, and *Bouteloua gracilis* (accession information in Supplemental table 1). Species were selected to represent different major lineages of the family *Poaceae* (Cynodonteae, Andropogoneae, Paniceae, Danthonieae, Poeae, and Paspaleae), and included both C_3_ (BEP and PACMAD clades) and a range of C_4_ species. In addition, we intentionally chose species within the same tribe that were previously reported to have varying responses (tolerant and sensitive) to low soil moisture. Seeds were germinated in 868.5 cm^3^ pots with a mix of potting soil and general-purpose sand with a ratio of 2:1 soil to sand and placed in a Kansas State University greenhouse under ambient conditions and raised to maturity throughout 2016–2018. After reaching maturity, the samples were subjected to 100% water reduction (referred to as ‘dry-down’), simulating an extreme drought, as previously described [62,63]. During the dry-down, samples were monitored daily and placed into categorical conditions based on their physiological state: “Initial”, “Stressed”, and “Recovery”. Physiological leaf traits were monitored daily and included: leaf-level net photosynthetic rates (*A*_n_; μmol CO_2_ m^-2^ s^-1^), stomatal conductance (*g*_s_; mol H_2_O m^-2^ s^-1^), transpiration (*E*: mmol H_2_O m^-2^ s^-1^), and instantaneous water use efficiency (*iWUE; A_n_/E*) calculated as the ratio between *A*_n_ and *E*. Data were collected with an LI-6400 system (LiCOR Biosciences Inc., Lincoln, NE, USA) equipped with an LED light source (light intensity maintained at 2000 μmol m^-2^s^-1^) CO_2_ concentration at 400 μmol mol^-1^, and relative humidity at ambient levels (35-50%). Physiological states were determined by relative rates of *A*_n_. The condition: “Initial” was measured on Day 1 (first day of drought after being watered the previous day) in order to avoid biased measurements from saturated soils. When samples reached near stomatal closure and extremely low photosynthetic rates (*A*_n_ < 25% of Day 1 *A_n_* (“Initial”)), they were categorized into the new condition “Stressed”. At this point, water was re-applied to soil saturation after the pertinent data were collected. Plants were allowed two days to recover before post-drought physiological data were collected (“Recovery”).

### Economic trait measurements

After physiological data were collected in the “Recovery” period, above and belowground tissues were harvested for all species. Leaf-level economic and anatomical data were collected from samples that included all non-droughted individuals but excluded *P. juergensii*, *P. notatum*, or *F. ovina* due to the lack of samples. The leaf tissue data included: leaf area (*LA*; cm^2^), specific leaf area (*SLA*, leaf area divided by dry mass; cm^2^ g^-1^), and leaf dry-matter content (*LDMC*, dry leaf mass divided by fresh mass; g g^-1^). *SLA* and *LDMC* were analyzed with the standardized rehydration method [56,64], while *LA* data were obtained by processing images in ImageJ [65]. Roots were washed and cleaned of debris for digital root imaging; analysis of root images was completed with a root imaging software (WinRhizo; Regent Instruments, Inc., Nepean, Ontario, Canada). Root imaging provided the following traits: total root length (cm), root diameter (mm), and specific root length (*SRL*, root length divided by dry mass; cm g^-1^). After scans were completed, above and belowground biomass samples were dried for 48 hours at 65°C and weighed for productivity comparisons.

### Anatomical trait measurements

The newest mature leaf was used for anatomical analysis prior to the initiation of drought from the following species: *Setaria viridis, Urochloa ruziziensis, Danthonia spicata, Rytidosperma semiannulare, Bouteloua dactyloides*, and *Bouteloua gracilis. Sorghastrum nutans, Andropogon gerardii*, and *Panicum virgatum* were collected from parent populations in the field at peak physiological performance and identical developmental stages. *Festuca ovina*, *Paspalum notatum*, and *Paspalum juergensii* samples were not included in these analyses due to sample loss. Leaf tissues for anatomical analyses, roughly 30 mm in length, were collected (4-8 samples per species; *n* = 33) by clipping leaf tissue and placing them into a fixative FAA (10% formalin / 5% glacial acetic acid / 50% ethanol (use 95% EtOH) / 35% DI water)) under a vacuum. Tissues were then cut (cross-sectioned) to 4μm in thickness with a Leica RM2135 microtome (Leica Biosystems, Newcastle, UK), and mounted in paraffin at Kansas State’s College of Veterinary Medicine Histopathology lab. Tissue was stained with Safranin-O and Fast Green [66], cover slipped, and imaged on a Zeiss 880 confocal microscope (Carl Zeiss, Walldorf, Germany) at 10X and 20X when necessary with a multitrack configuration, digital dual-bypass filters and a GaAsP detector (Fig. 1). Anatomical data were collected using IMAGEJ software [65] by analyzing two tissue regions from either side of the midrib between two major vascular bundles which were then averaged together from each leaf sample [58,59]. Here, the total subsampled area is referred to as the cross-sectional area or the area between two major vascular bundles (CSA). Anatomical traits collected from subsamples include: xylem area (*X*_area_; μm^-2^), xylem lumen diameter (*X*_diameter_; μm), *t/b* (xylem wall thickness/*X*_diameter_; xylem resistance to cavitation; μm μm^-1^); all xylem specific trait data were collected from major vascular bundles; while stomatal count (*S*_count_) was collected from the whole-leaf cross-section. In this study, we did not collect stomatal densities - as that would entail epithelial peels or impressions; therefore, we do not equate density measurements and interpretations with the *S*_count_. Instead, we utilized *S*_count_ to inform how many stomata are directly serving major and minor vascular bundles and other various photosynthetic tissues within the whole-leaf cross-section from which data were collected.

**Figure 1:**
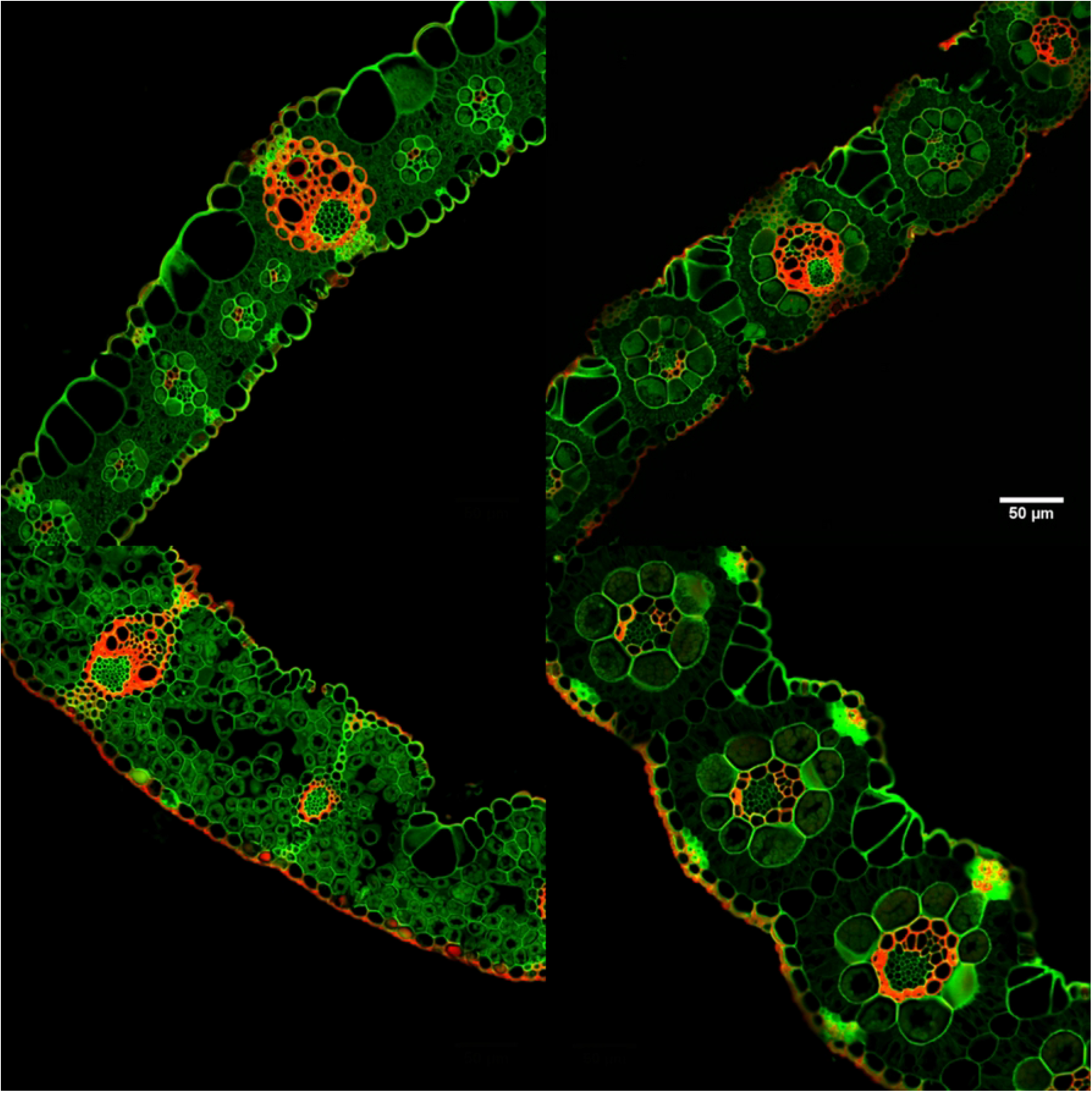
Leaf cross-sections of each major grass tribe stained with Safranin Red and Fast Green. Top left, *Andropogon* gerardii (Andropogoneae); top right, *Bouteloua dactyloides(Cynodonteae);* bottom left, *Danthonia spicata* (Danthonieae); bottom right, *Panicim virgatum* (Paniceae). Image taken with a Zeiss 880 confocal microscope.

### Data analysis

The selected traits were averaged by species and separated into the three physiological ‘stages’ (Initial, Stressed, and Recovery) based on physiological responses. We included tribe as a factor to investigate differences among lineages. We considered using a phylogenetic generalized linear mixed model (PGLMM), however, our goal was not to control for phylogeny, but rather to determine if lineages with different traits have evolved different drought tolerance strategies. All data were checked to meet assumptions of normality before analyses began. Comparisons among tribes and dry-down ‘stages’ were analyzed using mixed-effects analysis of variance (ANOVA) model with physiological data used as the response variables and tribe and condition as predictor variables (fixed effects). Tests were performed with the lmer function within the lme4 package in R [67]. To assess bivariate relationships between plant functional traits, we performed simple regression analyses (using the ‘lm’ function in R). Non-parametric data were analyzed via Kruskal-Wallis rank sum test paired with a post hoc pairwise Wilcox test. We also used Akaike’s information criterion, adjusted for small sample size AICc model selection to determine the most impactful trait parameters determining drought response using the “MuMIn” package in R [68,69]. All data were analyzed in the statistical program R V3.5.3 [70]. In order to summarize the relationships and range of physiological, functional, and anatomical diversity represented in our dry-down sample, we conducted a Principal Component Analysis (PCA) using the “prcomp” function within the “stats” library in R on the mean trait data across species, which cumulatively explained 72% of the variation in traits (Fig. 2). Not all traits were measured for every species, and so we focused on key traits coming from each of the data types we measured. The purpose of the PCA was to explore the multivariate relationships visually among species in multivariate space, and only include species that had all functional trait data (excluding *P. notatum*, *P. juergensii*, and *F. ovina*).

**Figure 2:**
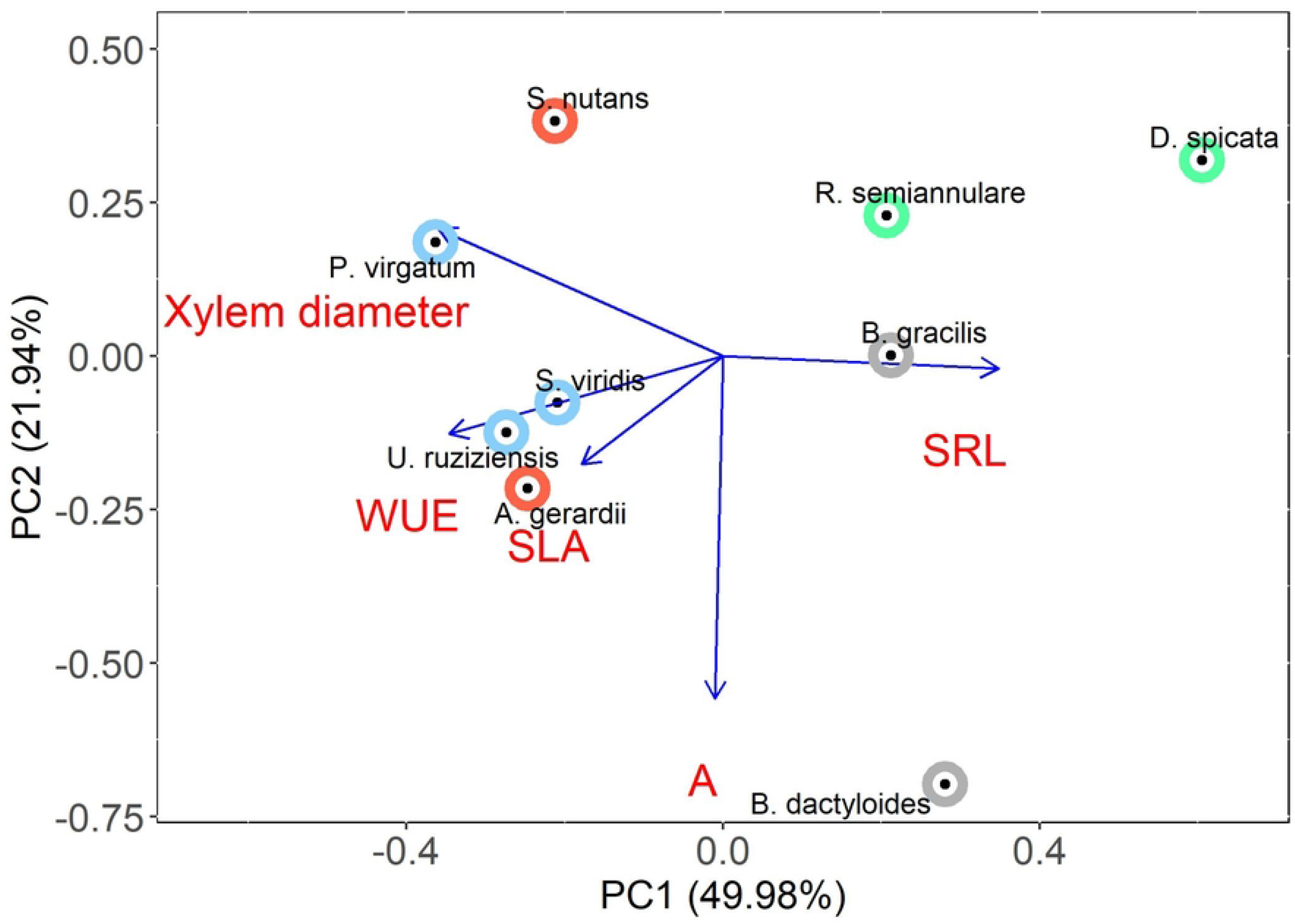
Principal components analysis (PCA) of mean trait values (in red text) of species in the dry down phase of the experiment. This PCA provides a summary of species in multivariate trait space using the first two PC axes, which together account for 72% of the trait variation. PC1 was most associated with variation in water use and rooting strategies whereas PC2 was primarily associated with photosynthetic rate. Information concerning PCA axes importance and subsequent loadings are located in Supporting Table 6. Andropogoneae (light red), Cynodonteae (grey), Danthonieae (green), and Paniceae (blue); each point is a species mean.

## RESULTS

Drought responses to the experimentally induced dry-down differed among species. Physiological viability, defined as maintaining at least 25% of the initial photosynthetic rate, ranged from 4-33 days (Fig. 3A). Drought duration (days in drought before re-watering) was similar among species within tribes, but varied significantly across tribes (Fig. 3A, *P* < 0.001). Species within the tribe Cynodonteae (*B. dactyloides* & *B. gracilis*) were physiologically viable for the longest period of time during the dry-down, reaching the “stressed” stage after 30 days with water withheld (Fig. 3A, Supporting Table S2), whereas the most drought-sensitive species were within Paspaleae and Poeae. These tribes were similar (*P* > 0.05), reaching the ‘stressed’ stage more than 20 days before the Cynodonteae tribe (Fig. 3A, Supporting Table S2). The recovery of grass species and tribes following re-watering displayed a more variable response (Fig. 3B). There were no statistical differences among tribes in their recovery dynamics (*P* > 0.05), though significant differences were observed among species and within tribes (*P* < 0.05). *S. nutans* and *B. gracilis* were the only species that exceeded pre-drought photosynthesis levels after recovery (114% and 121% of initial *A*_n_, respectively, Fig. 3B). *Festuca ovina* and *B. dactyloides* were the only species that did not regain at least 50% of ‘Initial’ physiological rates. Several species within Paspaleae, Paniceae, Danthoneae, and Andropogoneae lineages did not fully recover (100%) to ‘Initial’ physiological levels within the experimental timeframe. However, all recovered to at least 50% of Day 1 measurements (Fig. 3B). Andropogoneae species displayed the highest drought recovery, regaining on average over 92% of physiological functioning while the only species measured in Poeae (*F. ovina*) was the least resilient, with only 42% recovery of pre-drought *A*_n_ following re-watering.

**Figure 3:**
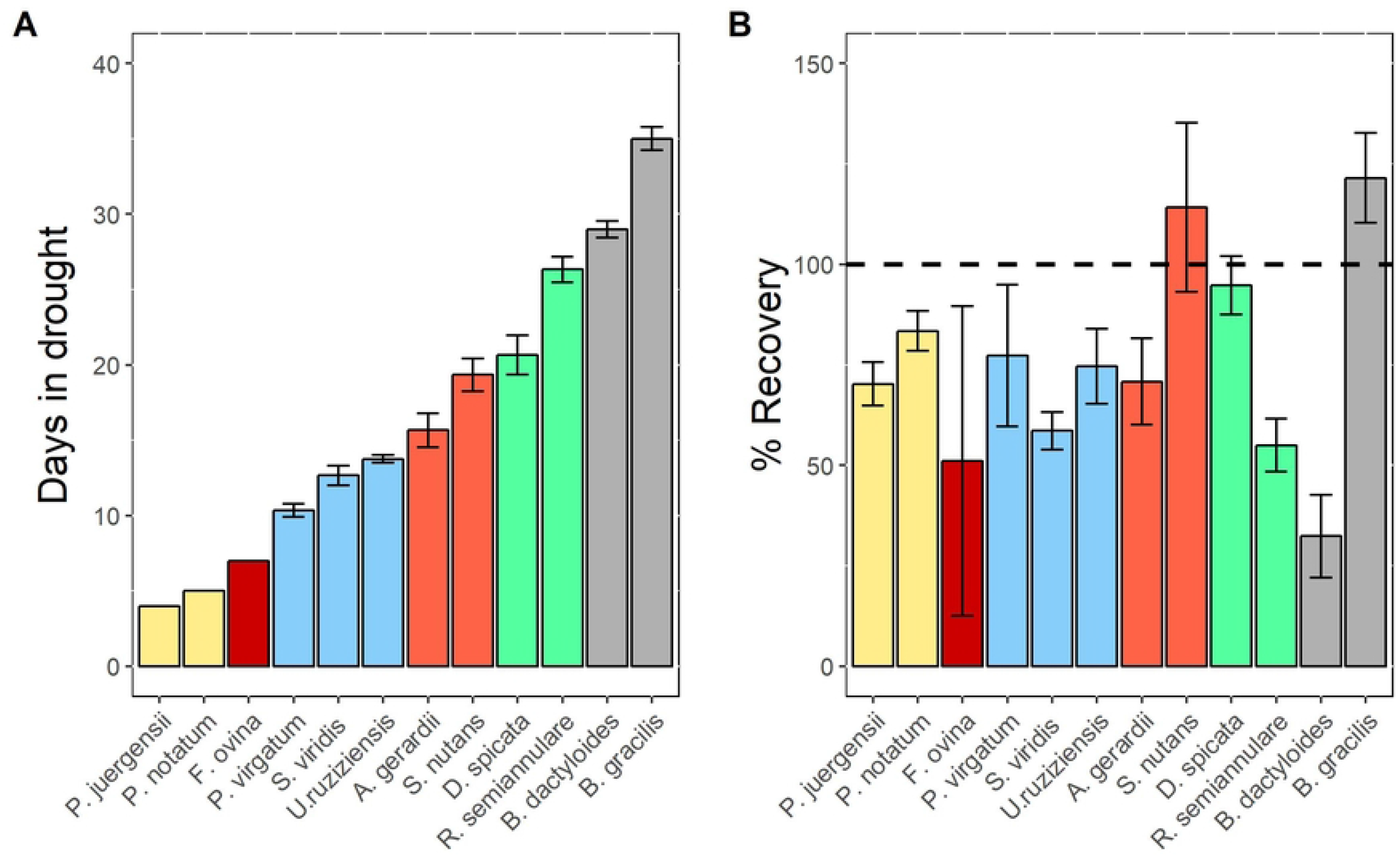
A) Number of days each species and tribe lasted before stomatal closure and rewatering occurred. B) The physiological recovery (A_n_) compared to Day 0 or Initial physiology (measured here as a percent). Dashed line signifies a complete 100% recovery of physiological function (i.e., A_n_ at or above its initial value). Andropogoneae (light red), Cynodonteae (grey), Danthonieae (green), Paniceae (blue), Paspaleae (yellow), Poeae (red); each point is a species mean and ± SE.

Leaf economic trait data were collected at the conclusion of the dry-down when recovery data were collected for each individual sample. Production of aboveground biomass was observed to vary significantly at both tribe and species level (Supporting Table S4, S5; *P* < 0.001). *SLA* was statistically similar within Andropogoneae, Cynodonteae, and Danthonieae (*P* > 0.05) while *SLA* within Paniceae species displayed significant variation (*P* < 0.05), ranging from 29 cm^2^ g^-1^ (*P. virgatum*) to 143 cm^2^ g^-1^ (*U. ruziziensis*). Similarly, *LDMC* was statistically similar across tribes, except for two species within Paniceae (*P* < 0.05) (*S. viridis* and *U. ruziziensis*) (Supporting Table S2). The production of fine root length (diameter < 0.5 mm) differed among tribe (*P* < 0.0001) and species (*P* < 0.0001). Significant differences in *SRL* were observed across tribes and species as well (*P* < 0.001, Supporting Table S2). All species within their respective tribes were found to have statistically similar *SRL* except for species in Paniceae (*P* < 0.05), Paspaleae (*P* < 0.05), and Danthonieae (*P* < 0.01) due to the 294.99 cm g^-1^ difference in *SRL*.

Most anatomical traits displayed significant differences among species across tribes (*P* < 0.05) but typically had reduced variability between species within the same tribe. *X*_area_ was statistically different between tribes (*P* < 0.001) with the exception of Danthonieae and Cynodonteae (*P* = 0.485). Andropogoneae (589.308 μm^2^) had the largest *X*_area_ and was five times larger than the smallest *X*_area_ found in Cynodonteae (108.957 μm^2^) (*P* < 0.01; Supporting Table S3). While there were significant species differences found across all tribes (*P* < 0.001; Supporting Table S3), there were no observable species differences within tribe (*P* > 0.05). *X*_diameter_ reflected a similar pattern to that of *X*_area_: significant differences between tribes (*P* < 0.001) and statistically similar values within tribes (*P* > 0.15; Supporting Table S3) (consistent with phylogenetic niche conservatism). Xylem resistance to cavitation (*t/b*) differed significantly across tribes (*P* < 0.001), species (*P* < 0.001), but not among species within a tribe (*P* > 0.05; Supporting Table S3). Stomata within the subsampled area (*S*_count_) showed high variation; the significant differences among tribes (*P* < 0.01) are likely attributed to Paniceae, which had higher *S*_count_ (Supporting Table S3).

There were few statistically significant relationships explaining drought responses and recovery among tribes (Supporting Fig. S1), except for *X*_area_ and *S*_count_. Surprisingly, given the large volume of literature designating *iWUE* as a pivotal functional trait reflecting drought tolerance, there was no statistically significant correlation of *iWUE* with drought resistance or resilience (*P* > 0.05). However, stomatal number was significantly negatively correlated with drought duration (*P* < 0.01) (Fig. 4), *X*_area_ (*P* < 0.05), and *iWUE* (*P* < 0.05) (Supporting Fig. S1). Results also indicate a differentiation between sample productivity and economic growth strategies (Fig. 5). A significant relationship was observed when comparing above and belowground biomass (Fig. 5A), yielding a tight positive relationship (*P* < 0.001; *R*^2^ = 0.825). Yet, when above and belowground economic strategies (*SLA* and *SRL*, respectively) were calculated, a breakdown in the previous relationship was observed (*P* > 0.05; *R*^2^ = 0.008) (Fig. 5B). While *SLA* displayed no bivariate relationships with other traits, *LDMC* correlated with *t/b* (*P* < 0.01) and *S*_count_ (*P* < 0.05) (Supporting Fig. S1). The AICc model selection process indicated how the selected functional traits influence both drought resistance and resilience. The model explaining the greatest variation in drought sensitivity included Tribe, *S*count, *iWUE*, *LDMC*, and *SLA*. However, the best explanation for variation in drought resilience included a single anatomical trait: *S*_count_.

**Figure 4.**
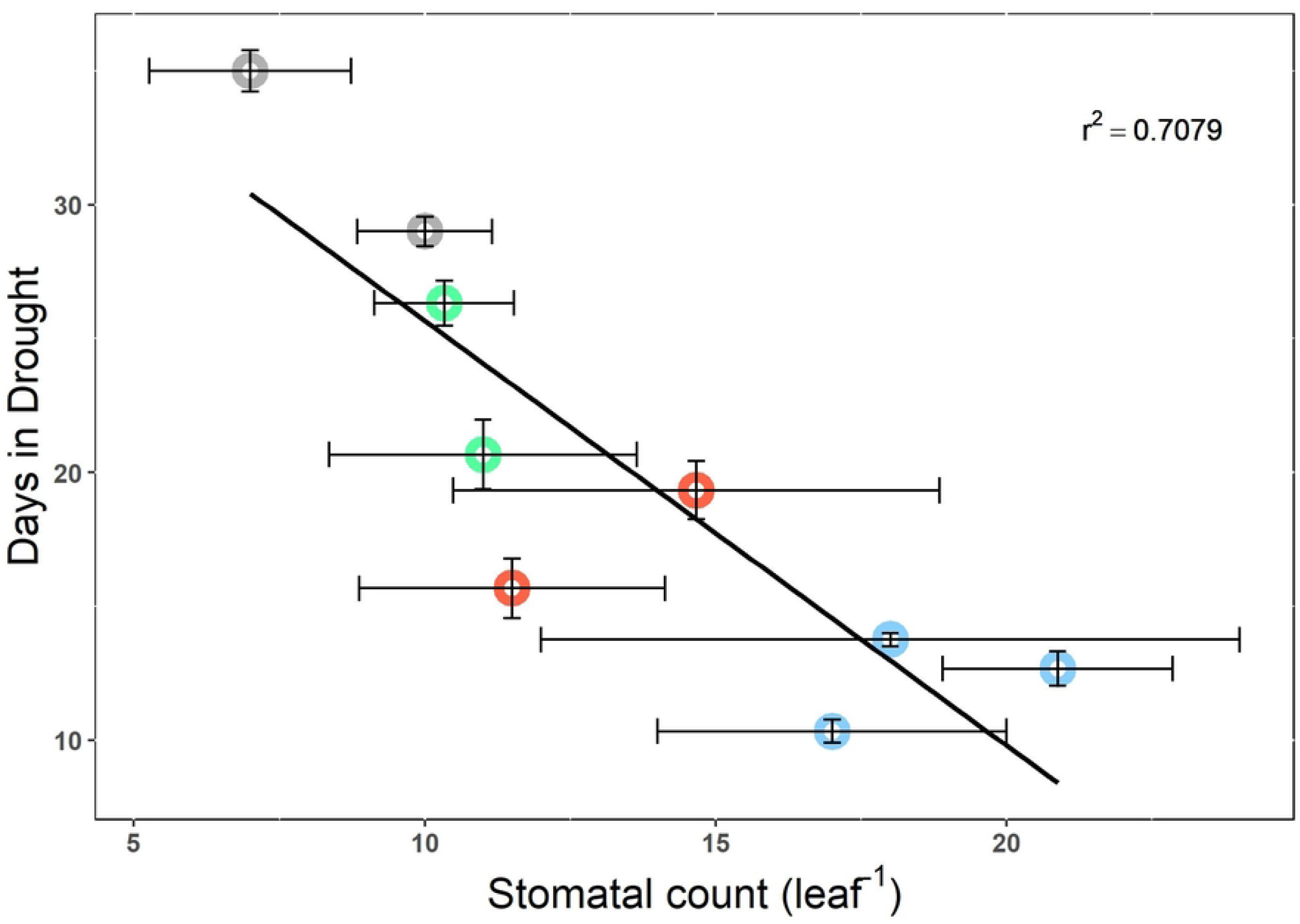
Relationship between stomatal count per entire leaf cross-section and days in drought before “Recovery”. Andropogoneae (light red), Cynodonteae (grey), Danthonieae (green), and Paniceae (blue); each point is a species mean and ± SE.

**Figure 5:**
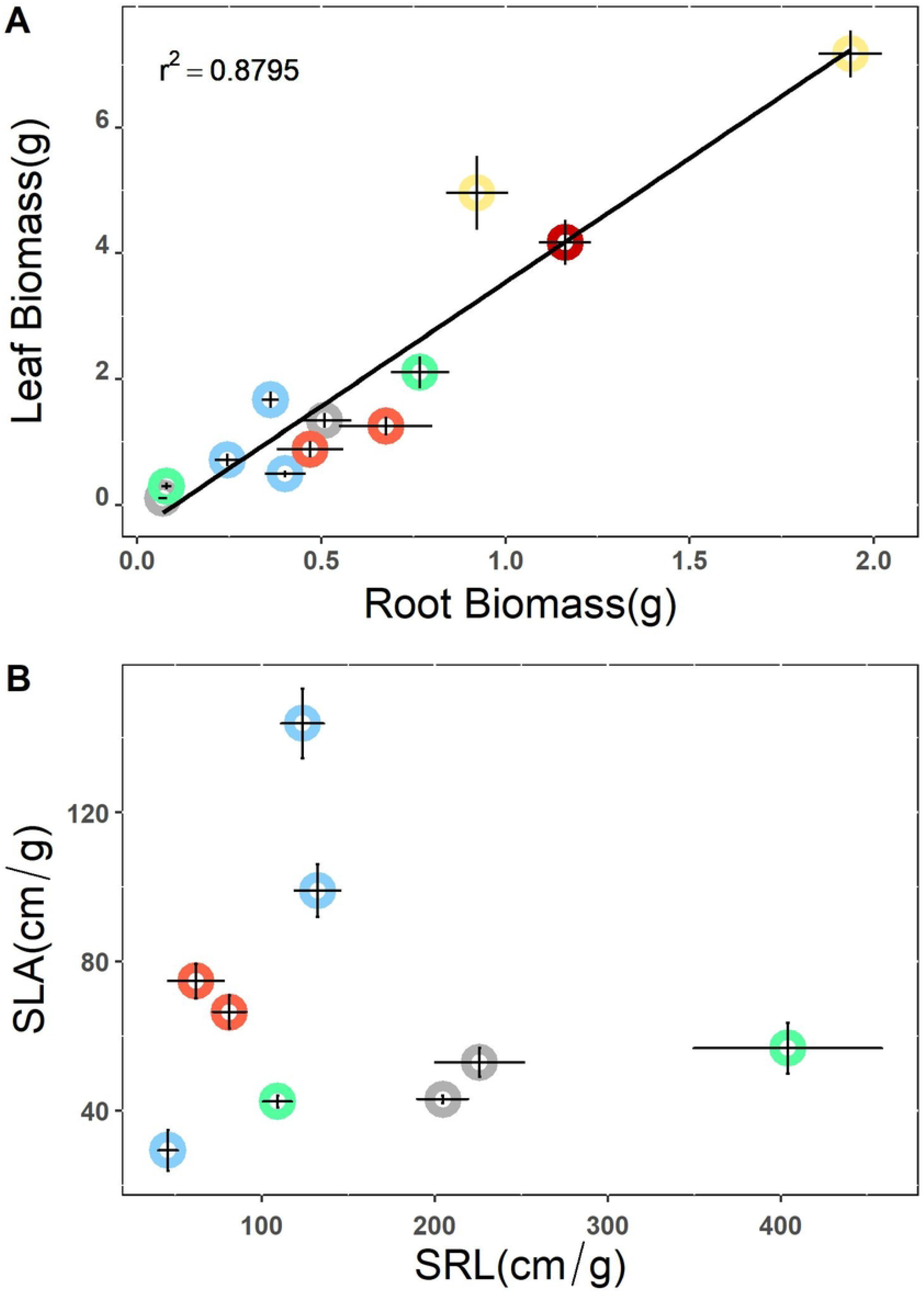
A) The relationship between leaf biomass and root biomass. B) Specific leaf area against specific root length. Andropogoneae (light red), Cynodonteae (grey), Danthonieae (green), Paniceae (blue), Paspaleae (yellow), Poeae (red); each point is a species mean and ± SE.

## DISCUSSION

Historically, water use efficiency (*WUE*) has been used as a seminal physiological trait to explain why some species persist and others succumb to drought [33,71–73]. However, the data presented here illustrate that using this physiological trait by itself may misconstrue the interpretation of drought responses in grass species (Fig. 3A; Supporting Fig. S1) [74]. Physiological, morphological (*SLA*, *SRL*, *LDMC*), and anatomical data (xylem area, stomata number, *t/b*) in combination help provide a more nuanced perspective on grass responses to drought, and when viewed as a collective, more appropriately identify mechanisms of water stress across diverse grass lineages. Shared phylogenetic and biogeographic histories have resulted in unique adaptations and trait development in Poaceae that are reflected in the patterns of global distribution and subsequent responses to drought [47,75,76]. Here, we show considerable variation in functional traits among diverse species within Poaceae in response to drought and recovery. In addition, these data illustrate that the traits and adaptations that confer an ability to withstand extreme drought conditions are not the same as the traits and adaptations that confer an ability to recover from drought, as they are driven by the coordination of different factors (Fig. 2; Fig. 3).

The grass species measured here displayed variable sensitivity to the dry-down, visible in both drought response and recovery (Fig. 3). The physiological responses observed during the dry-down were statistically related to variability in anatomical features, mainly those influencing water relations at the leaf level (i.e., Fig. 4). We also found differentiating relationships between productivity and economic strategies in both above and belowground tissues of the selected Poaceae species (Fig. 5), reflecting a trade-off in leaf and root growth economic strategies. Differences in economic traits indicated water-use strategies varied in drought responses [77–79]. For example, plants with lower *SLA* and higher *SRL* traits are likely to have lower metabolic costs and grow in resource-poor environments with an increased ability to acquire resources [56,80]. These trait strategies should allow for sustained physiological tolerance and quicker recovery following drought conditions. While evolutionary relatedness guides physiological, morphological, and anatomical traits in determining drought responses, drought recovery was mainly driven by *S*_count_ alone. The number of stomata within the subsampled cross-sectional area (CSA) is not equivalent to stomatal density measurement using epithelial peels. Rather, this proxy quantifies stomatal proximity as a vector. *S*_count_ indicates the number of stomata that are explicitly supplying CO_2_ to major and minor vascular bundles between two major vascular bundles. Stomatal density, rather, considers a leaf’s surface as a uniform and homogeneous surface and does not link stomatal and vascular bundle densities. Thus, *S*_count_ allows for a more direct mechanistic comparison with the other anatomical trait measurements performed here, (i.e., mesophyll area, bundle sheath area, and the diffusion distance through mesophyll) and may provide more detail on how stomata proximity to vasculature influences carbon assimilation and subsequent water loss [81–84].

Low soil moisture negatively impacts growth, increases xylem tension, and decreases carbon assimilation [85,86]. The ability to mitigate and recover from drought is based on anatomical and physiological traits [87,88]. While the impacts of severe drought on the physiology of grassland species have been observed in previous research, few studies combined physiological, whole-leaf, and anatomical trait data [47,89]. Additionally, our results indicate that closely related grasses can respond similarly to decreasing soil moisture (Fig. 3A; Fig. 4) but display variable responses when water becomes available (Fig. 3B). While we did not directly measure water availability in each pot, species were inherently variable in size (height, leaf size, etc.). Therefore, water was absorbed from the same volume of soil, lending to a greater understanding as to the species-specific responses. In addition, this variability supports previous claims that drought responses within a functional type (i.e., C_4_ grasses) are not uniform and vary due to a myriad of reasons (i.e., evolutionary histories, functional traits, etc.)[25,47]. The diversity in physiological responses (e.g., variability) among species has been observed to protect individuals and populations while protecting ecosystem functioning from the detrimental effects of drought [90–92].

Given the fundamental role that past evolutionary histories have played in shaping current species distributions [93,94], species that exhibit variable responses to ecosystem disturbances would benefit more than species that maintain static responses [95,96]. For instance, species that were more drought resistant (Cynodonteae) are broadly represented in the mixed and shortgrass prairies of North America, regions that are known to have less rainfall and more frequent drought [5]. In contrast, the drought-sensitive species (Paspalum) are from locations where soil moisture is typically not the most limiting resource [97]. In addition, Cynodonteae were also observed to have fewer stomata and decreased gas exchange rates compared to Paniceae and Paspalum species, leading to less water loss (Fig. 4; Supporting Table S2, S3). Therefore, it stands to reason that phylogenetically dissimilar species evolving under different environmental constraints would exhibit disparate drought response to the imposed dry-down, while more closely related species would respond more uniformly (Fig. 3A; Fig. 4). Our data also display a clear indication of an evolved plasticity in physiological responses to variable climate conditions, as native grasses typically occupy regions with similar climate variability [98]. Grasses sampled in this experiment were severely desiccated and recovered >50% of pre-drought physiological functioning, and in several cases, physiological rates that were 20–30% higher than the initial state (Fig. 3B), highlighting a potentially unique characteristic of grasses across lineages. The ability to acquire water and other nutrients quickly following drought disturbances likely facilitates grasses competing with other neighboring functional types with deeper access to water [99,100].

Water availability directly impacts plant physiological responses, which are constrained by internal anatomical machinery (Fig. 1) [58,101]. For example, the spatial separation of C_4_ photosynthesis allows for a reduced stomatal conductance and decreased water loss, leading to higher water-use efficiencies [102–105]. Our findings, however, do not support this claim (Supporting Fig. S1). *iWUE* was not observed to directly aid in drought sensitivity or recovery of grasses in this study, but it was positively related to the number of stomata (Supporting Fig. S1), indicating here, that the presence of more stomata is associated with higher *iWUE*. This counterintuitive result does not indicate a more drought tolerant strategy; rather, it suggests higher gas exchange rates resulting in greater water loss which leads to quicker desiccation [12,38,106]. However, previous research has indicated that stomatal patterning, morphology, and densities can greatly influence/alter physiological responses to water stress [107,108]. Figure 4 clearly indicates species with more stomata have an increased sensitivity (decreased resistance) to drought, which may require reevaluations of previously held claims regarding the functional significance of *WUE* [109].

Xylem characteristics have also been previously shown to impact an individual’s water–use [88,110,111]. Xylem area is a commonly measured trait because it corresponds with the amount of water that can be transported at any given time. Here, our results indicate two water transport strategies. Larger xylem (*X*_area_) decreases drought resistance while displaying a positive relationship with recovery, when excluding *A. gerardii* and *S. nutans* (Supporting Fig. S1). This strategy enables individuals with larger *X*_area_ to transport greater amounts of water, when available (seen in recovery). But, drought conditions can lead to increased tension on the water column inside xylem vessels, ultimately increasing the potential for embolism formation during drought conditions [112]. Previous research has highlighted how increased thickness of xylem wall tissue with smaller diameter lumen (*t/b*) can protect from embolism events in water limiting conditions [113,114]; however, our data do not corroborate such findings (Supporting Fig. S1). Anatomical traits were observed to contain large variation, which we can contribute to two main factors: (1) our sample size was relatively small, due to the time - consuming nature of anatomical studies; and (2) anatomical traits are complex in nature and have large variability among individuals and within grass leaves (Fig. 1) [58,82].

While anatomical traits and leaf-level physiological rates provide key mechanistic insights into drought sensitivity and resilience, whole-plant traits are more easily observable and require less detailed scientific instrumentation and training [57,77]. Whole-plant traits illustrate broader growth strategies by the individual, such as resistance or avoidance of detrimental growth conditions. Our results indicate a linear relationship between above and belowground productivity (Fig. 5A), indicating a constant proportional investment by the selected grasses. However, when comparing two widely utilized traits within the leaf and root economic spectrum (*SLA* and *SRL*), the previous relationship breaks down to reveal tradeoffs in grass growth strategies (Fig. 5B). Individuals that invest in a root system designed for quick absorption of water and nutrients (high *SRL*) may produce an inexpensive leaf (low *SLA*), while more ‘expensive’ leaves (high *SLA*) appear to be associated with a less economically efficient rooting strategy (Fig. 5A). This finding highlights the inability of grasses to produce tissues at the upper limits of the leaf and root economic spectrum. However, it’s important to recognize and consider the potential for convoluted interpretations of economic traits, as they are the amalgamation of several underlying traits [56,58]. In addition, there could be grasses, which were not investigated here, that invest in high *SLA* and *SRL*. One of these complex traits is leaf dry matter content (*LDMC*), which was correlated with the thickening of xylem vessel walls or xylem reinforcement: *t/b*. This thickening of water transport tissues (or xylem reinforcement) increases the strength at which the water column can withstand tension, allowing for a more negative water potential while decreasing the likelihood of cavitation, a physiological indicator of drought stress [112].

There is currently a dearth of available functional trait data in grasses, an underrepresented functional type in trait databases given the importance of grass species for food and forage and their extensive geographical coverage. Results from this study indicate the need for increased collection of grass functional traits across a diverse assemblage of species within a functional type. Plant functional types are often used in ecosystem models to more easily group plants by common features; however, as our results indicate, this may lead to poor parameterization and model output as such functional types do not account for phylogenetic relatedness. Our AICc selection indicates that anatomical traits (specifically stomatal count), physiological, economic traits, and phylogeny were essential to understanding species ability to withstand drought, while stomatal count was the best explanation for recovery responses. The trait data from these lineage-specific responses to drought have potential consequences for how different grasslands are represented and forecast in Earth System Models [25,115].

The evolutionary histories of lineages within Poaceae have led to the development of unique morphology (leaves and roots), anatomy (stomatal shape and photosynthetic tissue arrangement), and physiology; these traits have allowed this functional type to dominate much of the surface cover on every continent (excluding Antarctica). Phylogenetic divergences and subsequent trait adaptations have led to contrasting responses and recovery from drought (Fig. 3). Anatomical traits were key in explaining physiological drought response and recovery, specifically traits concerned with water usage. Surprisingly, species that exhibited increased *iWUE* were more prone to quicker desiccation, which is most likely due to the same individuals maintaining larger numbers of stomata and higher overall rates of gas exchange (Supporting Fig. S1). Interestingly, this same trait (*S*_count_) was responsible for recovery from drought, which did allow for faster drought recovery. This study underscores the importance of collecting a myriad of in-depth trait data from several Poaceae lineages to understand better the mechanisms that describe drought responses and recovery.

## ACKNOWLDGEMENTS

We thank the Konza Prairie LTER program (NSF DEB-1440484), the NSF Dimensions of Biodiversity program (NSF DEB 1342787), and the NSF Macrosystems program (NSF 192635). We also thank Ryan Estes and Sam Sharpe for their help in collecting gas exchange data on several occasions, and Joel Sanneman at Kansas State Universities’ confocal core facility for his help and expertise in attaining confocal images. The findings and conclusions are those of the authors and should not be construed to represent any official USDA or U.S. Government determination or policy.

## SUPPLEMENTARY DATA

Fig S1. Correlation matrix of bivariate physiological, morphological, and anatomical traits for each species.

Table S1. Source and accession numbers for all species with each associated subfamily and tribe.

Table S2. Mean physiological data and standard error for all species at each dry down stage.

Table S3. Mean anatomical data and standard error for all species. NA refers to species that have no data.

Table S4. Mean leaf – based morphological data for each species with standard error (SE). NA refers to species that have no data.

Table S5. Mean root – based morphological data for each species with standard error (SE). NA refers to species that have no data.

Table S6. Summary statistics and variance measurements from PCA axes and PCA loadings of the traits for the top three PCA axes from Figure 2.

## AUTHOR CONRIBUTIONS

SB, MZ, MU and JN planned and executed data collection. All authors contributed to the writing of the manuscript. DG, CS, JN and MU selected the species.

## FUNDING

Konza Prairie LTER program (NSF DEB-1440484)

NSF Dimensions of Biodiversity program (NSF DEB 1342787)

NSF Macrosystems program (NSF 192635)

## DATA AVAILABILITY

The datasets used and/or analyzed during the current study are available from the corresponding author on reasonable request. In addition, mean data is available in the supplementary materials.

